# Synthetic model ecosystem of 12 cryopreservable microbial species allowing for a noninvasive approach

**DOI:** 10.1101/2020.10.23.351742

**Authors:** Kazufumi Hosoda, Shigeto Seno, Naomi Murakami, Hideo Matsuda, Yutaka Osada, Rikuto Kamiura, Michio Kondoh

**Affiliations:** RIKEN Center for Biosystems Dynamics Research, 6-2-3 Furuedai, Suita, Osaka, 565-0874, Japan; Center for Information and Neural Networks (CiNet), National Institute of Information and Communications Technology (NICT), Osaka, Japan; Institute for Transdisciplinary Graduate Degree Programs, Osaka University, 1-5 Yamadaoka, Suita, Osaka 565-0871, Japan; Life and Medical Sciences Area, Health Sciences Discipline, Kobe University, Tomogaoka 7-10-2, Suma-ku, Kobe, Hyogo, 654-0142, Japan; Graduate School of Information Science and Technology, Osaka University, 1-5 Yamadaoka, Suita, Osaka 565-0871, Japan; Graduate School of Life Sciences, Tohoku University, 6-3 Aoba, Aramaki, Aoba-ku, Sendai 980-8578, Japan

**Keywords:** Synthetic ecosystem, Experimental model ecosystem, Microbial microcosm, Machine learning

## Abstract

Simultaneous understanding of both individual and ecosystem dynamics is crucial in an era marked by the degradation of ecosystem services. Herein, we present a high-throughput synthetic microcosm system comprising 12 functionally and phylogenetically diverse microbial species. These species are axenically culturable, cryopreservable, and can be measured noninvasively via microscopy, aided by machine learning. This system includes prokaryotic and eukaryotic producers and decomposers, and eukaryotic consumers to ensure functional redundancy. Our model system displayed both positive and negative interspecific interactions and higher-order interactions that surpassed the scope of any two-species interaction. Although complete species coexistence was not our primary objective, we identified several conditions under which at least one species from the producers, consumers, and decomposers groups, and one functionally redundant species, persisted for over six months. These conditions set the stage for detailed investigations in the future. Given its designability and experimental replicability, our model ecosystem offers a promising platform for deeper insights into both individual and ecosystem dynamics, including evolution and species interactions.

## Introduction

Understanding ecosystem dynamics has become increasingly crucial in light of the recent degradation of ecosystem services (Brondizio et al., 2019; Duraiappah et al., 2005). Since all species in ecosystems undergo persistence and fluctuations as a consequence of interactions with various other species, the conservation and management require an understanding of the dynamics of not only individual species, but also the entire ecosystem (Sinclair and Byrom, 2006). Fundamentally, individual species and entire ecosystems are intrinsically interdependent in hierarchical biological systems, necessitating the simultaneous consideration of both their dynamics (Conrad, 1996; Conrad and Pattee, 1970; Hosoda et al., 2016; Nakajima, 2021). While the dynamics of individual species have been extensively studied using “model organisms” (Fields and Johnston, 2005; Hale et al., 2003) such as *Escherichia coli*, experimental investigations of entire ecosystems present challenges. By constructing controlled, replicable ecosystems, researchers can dissect multifaceted interactions among system components and predict potential cascade effects of disturbances (Beyers and Odum, 1993; Odum and Barrett, 2005). However, compared with the organism level, experimental ecosystems have intrinsic difficulties in handling, replicability, and experimental commonality. If there is a “model ecosystem” that can be handled as easily as *E. coli* experiments, the understanding of dynamics of both individual populations and entire ecosystems will be greatly accelerated. In this study, we developed a high-throughput experimental system of species-defined microbial synthetic ecosystems that are noninvasively measurable.

Species-defined microbial experimental ecosystems, known as gnotobiotic microcosms (Beyers and Odum, 1993; Nixon, 1969; Taub, 1969), have been recognized as a type of experimental model ecosystems (Widder et al., 2016). As a model for understanding community dynamics, all species should be identified, because it is well-known that the existence of rare species can have a large impact on the entire ecosystem (Mouillot et al., 2013; Paine, 1969). Furthermore, because it is an experimental system for microorganisms, it is easy to handle owing to its small size. Although properties unique to large-scale ecosystems cannot be studied, gnotobiotic microcosms are useful for understanding universal properties common to other ecosystems, just as the model organism *E. coli* is useful for a wide range of other organisms (Blount, 2015; Zimmer, 2012). Specifically, gnotobiotic microcosms were used not only for understanding the effect of species composition on stability (Tanaka et al., 1994), community assembly (Drake, 1991; Fukami, 2015), trophic structure (Naeem and Li, 1998), and dynamics of complex communities in microbiota (Chodkowski and Shade, 2017; Grosskopf and Soyer, 2014; Ponomarova and Patil, 2015; Tan et al., 2015; Widder et al., 2016), but also for the application in bioprocessing (Ben Said and Or, 2017; Brenner et al., 2008; Shong et al., 2012) and to environmental problems such as the effect of chemicals or genetically-modified organisms (Benton et al., 2007; Fuma et al., 2005; Inamori, 2020).

Among gnotobiotic microcosms, we designate synthetic assemblages of microorganisms that are axenically culturable and cryopreservable as SCMs (synthetic cryopreservable microcosms). Despite the drawback of not perfectly mirroring specific natural ecosystems, SCMs are regarded as ideal model ecosystems for several reasons (Karkaria et al., 2021; Kawabata et al., 1995; Mee and Wang, 2012; Momeni et al., 2011; Shou et al., 2007; Taub, 1969). First, because the component organisms are cryopreservable, the experimental conditions that include the state of the organisms are replicable, and it is possible to compare a species before and after evolution. Evolutionary adaptation is crucial for ecosystem dynamics and multispecies coexistence (Kondoh, 2003; Thompson, 1998) (Schoener, 2011; Toju et al., 2017), and thus it is important for model ecosystems that component organisms are cryopreservable. Secondly, SCMs provide complete design control over both biotic and abiotic constituents, unlike microcosms that are composed of species isolated from nature and remixed (Hu et al., 2022; Tanaka et al., 1994). For instance, the use of model organisms can minimize unknown elements associated with each species and ensure the absence of unintended symbiotic pairs. Specifically, SCMs have been used to understand evolution (Azuma et al., 2023; Momeni et al., 2011; Nakajima et al., 2009; Waite and Shou, 2012), historical contingencies in succession (Chuang et al., 2019; Hekstra and Leibler, 2012), and higher-order microbial interactions (Mickalide and Kuehn, 2019).

However, existing configurations of SCMs often suffer from shortcomings in terms of ecological diversity. First, many SCMs lack predatory components (Hosoda et al., 2011; Momeni et al., 2011), thus omitting crucial trophic interactions that can drive ecosystem stability and complexity. This absence can be critical because trophic interactions are fundamental to our understanding of ecological dynamics (Blasius et al., 2020; Paine, 1966; Ripple et al., 2016; Terborgh et al., 2001). Second, in many SCMs, one of the functional groups, such as producers, consumers, or decomposers, is represented by a single species (Frentz et al., 2015; Kawabata et al., 1995; Nakajima et al., 2009; Taub, 1969). This limitation curtails the exploration of redundancy and competitive interactions, a central factor in community dynamics (Gauze, 1934; May, 1972; Mougi and Kondoh, 2012; Naeem, 1998). The absence of functionally redundant species also makes it difficult to distinguish between the effects of functional groups and species identity. Addressing these gaps in predation and functionally redundant species is important for model ecosystems to mimic and provide insights into their natural counterparts.

In this study, we developed a high-throughput experimental system of SCM consisting of 12 functionally and phylogenetically diverse species with known genomes that are axenically culturable, cryopreservable, and noninvasively measurable by microscopy using a machine learning model. The 12 species of microorganisms include more than one species of prokaryotic and eukaryotic producers and decomposers, and eukaryotic predators as consumers, that is, they are functionally redundant. These species exhibited both positive and negative interspecific interactions in paired cultures, and demonstrated higher-order interactions in microcosm experiments that were not accounted for by any single two-species interaction. Note that the coexistence of all 12 species is not the goal here, because competitive exclusion itself is important for understanding ecosystem dynamics (Gauze, 1934). However, because a certain degree of coexistence is necessary as a foothold for future research, we investigated the conditions where at least one species from each of the three functional groups, in addition to one redundant species, survived. We show various conditions under which such diverse species survived for longer than six months in this SCM.

## Results and Discussion

### Basic design of the model ecosystem

The constituent organisms were 12 diverse species of microorganisms, as shown in Table 1 (see Materials and Methods for preparation). These 12 species included three important functional groups in ecosystems: producers, consumers, and decomposers. We here refer to the non-decomposer heterotrophs as consumers, which are both protozoans. Except consumers, each functional group comprises multiple species from both eukaryotic and prokaryotic domains. All of these species have been well-studied; they can be axenically cultured and cryopreserved, and their genomes are known. In addition, the 12 species include motile organisms such as *E. coli*, *Dictyostelium discoideum*, *Tetrahymena thermophila*, and *Chlamydomonas reinhardtii*, and organisms with multicellular structures such as *Candida parapsilosis* and *Anabaenopsis circularis*; thus, this SCM can be used for research including spatial structure in the future.

**Table 1.** Species used.

|  |  | Species | Classification | Strain | Shape | Fluorescence |
| --- | --- | --- | --- | --- | --- | --- |
| Decomposers | #0 | *Escherichia coli* | Proteobacteria | DH1ΔilvE:: (dsred.T3-cat)<sup>a</sup> | Rod | ex.540/em.580 |
|  | #1 | *Staphylococcus epidermidis* | Firmicutes | JCM2414<sup>b</sup> | Spherical | (-) |
|  | #2 | *Schizosaccharomyces pombe* | Ascomycota | L972<sup>c</sup> | Rod | (-) |
|  | #3 | *Saccharomyces cerevisiae* | Ascomycota | BY25611<sup>d</sup> | Oval | ex.485/em.515 |
|  | #4 | *Candida parapsilosis* | Ascomycota | JCM1612<sup>b</sup> | Oval/Filamentous | (-) |
| Consumers | #5 | *Dictyostelium discoideum* | Amoebozoa | AX2<sup>e</sup> | Amoeba | (-) |
|  | #6 | *Tetrahymena thermophila* | Ciliophora | CU427&CU428<sup>f</sup> | Oval | (-) |
| Producers | #7 | *Anabaenopsis circularis* | Cyanobacteria | NIES21<sup>g</sup> | Filamentous | ex.615/em.655 |
|  | #8 | *Synechocystis* sp. | Cyanobacteria | PCC6803<sup>h</sup> | Spherical | ex.615/em.655 |
|  | #9 | *Raphidocelis subcapitata* | Chlorophyta | NIES35<sup>g</sup> | Crescent | ex.480/em.685 |
|  | #10 | *Chlorella vulgaris* | Chlorophyta | NIES227<sup>g</sup> | Spherical | ex.480/em.685 |
|  | #11 | *Chlamydomonas reinhardtii* | Chlorophyta | NIES2236<sup>g</sup> | Oval | ex.480/em.685 |

(a) Constructed from DH1 (Hosoda et al., 2011)
(b) Provided by Japan Collection of Microorganisms (JCM), RIKEN BRC, Japan
(c) A kind gift from Dr. Kojiro Ishii of Kochi University of Technology
(d) BY4741ho::Kan-TEFIlpr-GFP-ADH1ter (Breslow et al., 2008), provided by the National Bio-Resource Project (NBRP), Japan
(e) DBS0235521, provided by the Dicty Stock Center, Northwestern University, USA
(f) Conjugated progeny of CU427 and CU428, provided by the Tetrahymena Stock Center, Cornell University
(g) Provided by the NIES collection, Japan
(h) ATCC27184, provided by the American Type Culture Collection (ATCC), USA

These species can be distinguished not only by methods such as PCR, which detects genomic differences, but also noninvasively by their shape and fluorescence. Fluorescence spectroscopy can measure the amount of fluorescent-labeled *E. coli*, fluorescent-labeled *Saccharomyces cerevisiae*, cyanobacteria (autofluorescence), and green algae (Chlorophyta; autofluorescence), although it is not possible to distinguish the species within the cyanobacteria or within the green algae. Microscopy can identify all these species, but only within a limited area near the bottom surface.

As a container, a black 384-well plate was used to make it a high-throughput system. Since most organisms exist near the bottom due to gravity, light was illuminated from the bottom (Figure 1). We used a liquid medium based on half the concentration of BG-11 medium, a standard medium for cyanobacteria (Allen, 1968), supplemented with 1% LB medium, a standard medium for bacteria (Bertani, 1951) (see Materials and Methods for details).

**Figure 1.**
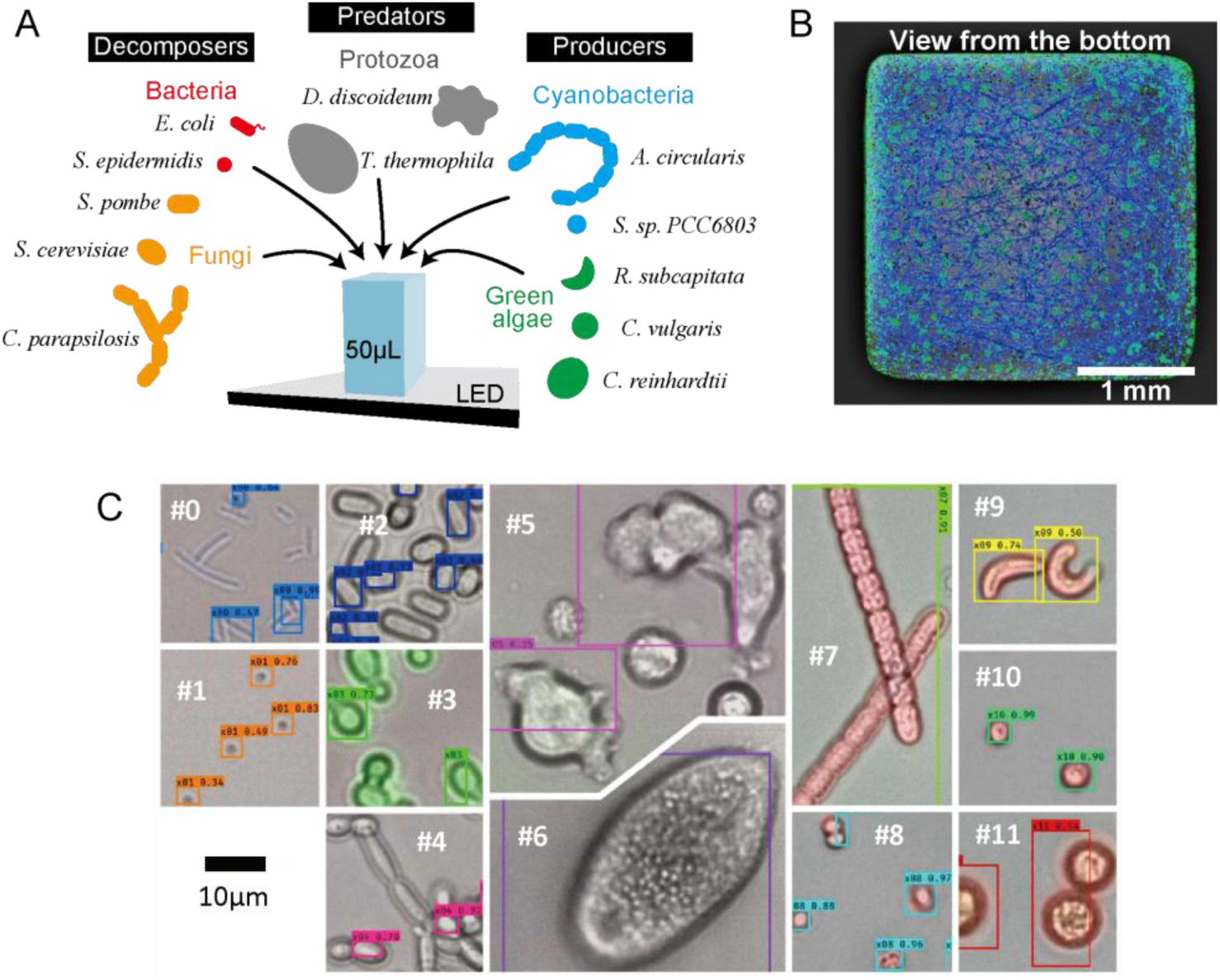
Schematic of the synthetic cryopreservable microcosms. (A) The container 384-well plate (a well of the plate is depicted) was placed on the LED panel. (B) An example of a microscopic image taken from the bottom. Fluorescent images of *E. coli* (red, but difficult to observe), cyanobacteria (blue), and green algae (green) were overlaid on the bright-field image. (C) Examples of micrographs of each microorganism and their detection using machine learning.

### Detection by machine learning

A prominent feature of this experimental system is that the populations of all 12 species are noninvasively measurable by identifying each individual microorganism in micrographs of microcosms using a machine learning model, in addition to detection by fluorescence spectroscopy. Noninvasive measurements have several advantages: (i) applicability to completely closed systems, (ii) high-throughput capacity by omitting sample collection, and (iii) avoidance of system-volume reduction and/or population disturbance due to sample collection. In this system, fluorescence spectroscopy enables us to measure populations with specific fluorescence properties, *e.g.*, green algae, but it is not possible to know the species within green algae. Therefore, microscopy with machine learning was used to complement it, although its accuracy was lower than that of fluorescence spectroscopy.

Detection of microorganisms, including bacteria and larger plankton, by microscopy with machine learning has been well-studied (Kosov et al., 2018; Kulwa et al., 2019; Kumar and Mittal, 2010; Lumini et al., 2020; Panigrahi et al., 2020), and we used an object detection network framework YOLOv3 (Redmon and Farhadi, 2018) (see Materials and Methods for details) as a conventional fast and accurate system for detecting objects in images.

Using this method, 12 species were detected, as shown in Figure 2A (detection examples are shown in Figure 1C). The misidentification errors were less than 1% for nine of the twelve species (except #4,5,6 in Table 1). Errors greater than 1% for both protozoans (#5 and #6) were mainly due to the false detection of tiny particles excreted from protozoan cells as cocci (#1; see Figure S3). This misidentification will be avoided in the future by fluorescently labeling cocci. In any case, both protozoa were rarely dense enough for these particles to be detected, and were rarely erroneously detected as cocci in the microcosm experiments.

**Figure 2.**
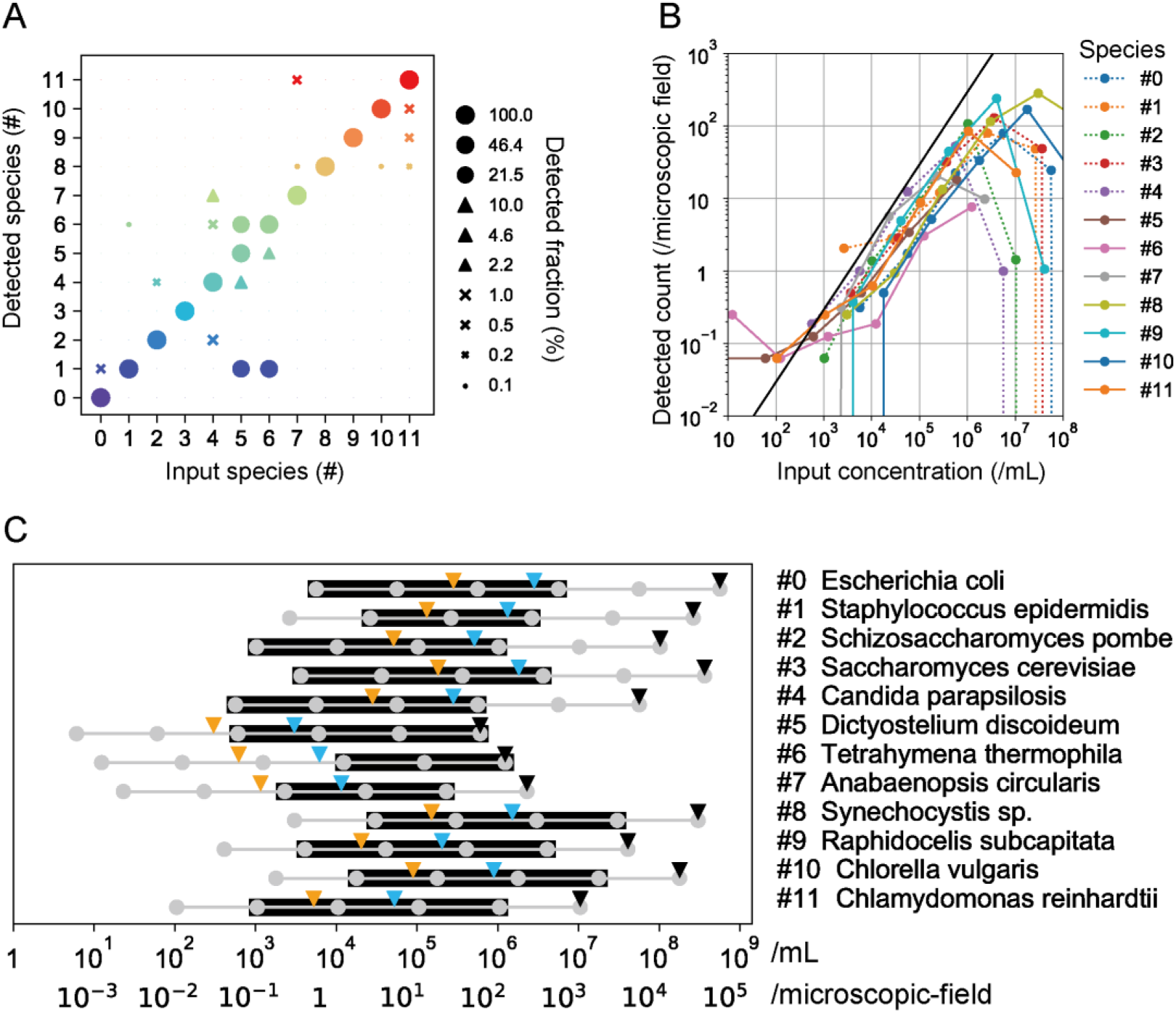
Detection of each species using a machine learning model. Individual microorganisms in images of monocultures (different from that used for training) were detected by the machine learning model using YOLOv3 (details are shown in Text S1 and Figure S2). (A) Detection accuracy. The fractions of the detected species are shown on the vertical axis when we tested monocultures of species on the horizontal axis. We used the data of 36 replicated wells from 9 independent plates for a single species. (B) Calibration curves for each of the 12 species. The mean value of 16 replicated wells from 4 independent plates is plotted. The black line represents the value when the input microorganisms are counted. Details, such as deviation for each plot, are shown in supplementary data (Figure S2). Note that species #4 and #7 exhibit filamentous structures. Species #4 was counted on a per-cell basis because it could also exist as a unicellular yeast and the training label was created on a per-cell basis. Species #7 was quantified as a multicellular organism. In both cases, there is a significant variation in the number of cells within a single cluster, leading to a greater variability when compared to the other species. Although the black lines were based on counts from visual inspection, the height of the lines should also include greater errors for the same reasons. (C) Concentration ranges of monocultures of each species. Black triangles show the saturation concentration in monocultures. Blue and orange triangles show the high and low initial concentration used in the microcosm experiments, respectively. Gray circles show the concentrations of the sample prepared by diluting the saturated monocultures, and those diluted samples were used for the standard curves in (B), and black lines show the ranges where the detected counts correlate with the input concentration (see Figure S2 for detail).

Figure 2B shows the standard curve for each species. We found a range wherein the counted amount correlated with the input amount for each of all species. These ranges were approximately 10^4^–10^6^/mL (summarized in Figure 2C; 500–50,000/well or 1– 100/microscopic field; specific ranges for each species are shown in Figure S2), which could be applied in the semi-open microcosm experiments shown below (except for protozoa). Unlike most bioanalytical methods, the lower detection limit mainly depended on the presence of microorganisms in the micrograph, as the background at the blank was rarely observed for nine species (other than #1, #5, and #6; see Figures S2B and S3). The detection efficiency (difference from the black line in Figure 2B) within these range was greater than 10% for most of the species. The efficiency depends not only on machine learning but also on the fraction of microorganisms present at the center of the well bottom (see Figure S2).

At high concentrations, the detected counts decreased steeply because the microorganisms overlapped in three dimensions, and were therefore indistinguishable. This occurs not only in the case of one species, but also in the case of a mixture of multiple species, that is, high concentrations of one species make it impossible to detect the other species (see Figure S4A and S5, for example micrographs and interfered counts, respectively). This high-concentration inapplicability is a limitation of this method, which is not due to machine learning, because this also happens to the human eye. In addition, the morphology of an organism itself may change depending on the culture conditions and passages, which may lead to detection errors. For instance, during the ecological experiments, species #10 tended to increase in size, which often led the machine learning algorithm to misidentify it as species #11 (e.g., Figure 3D). Solving these problems is a future challenge. However, there remain issues, but this is a great advantage in that it can be detected with a certain accuracy in a high-throughput and non-subjective manner.

**Figure 3.**
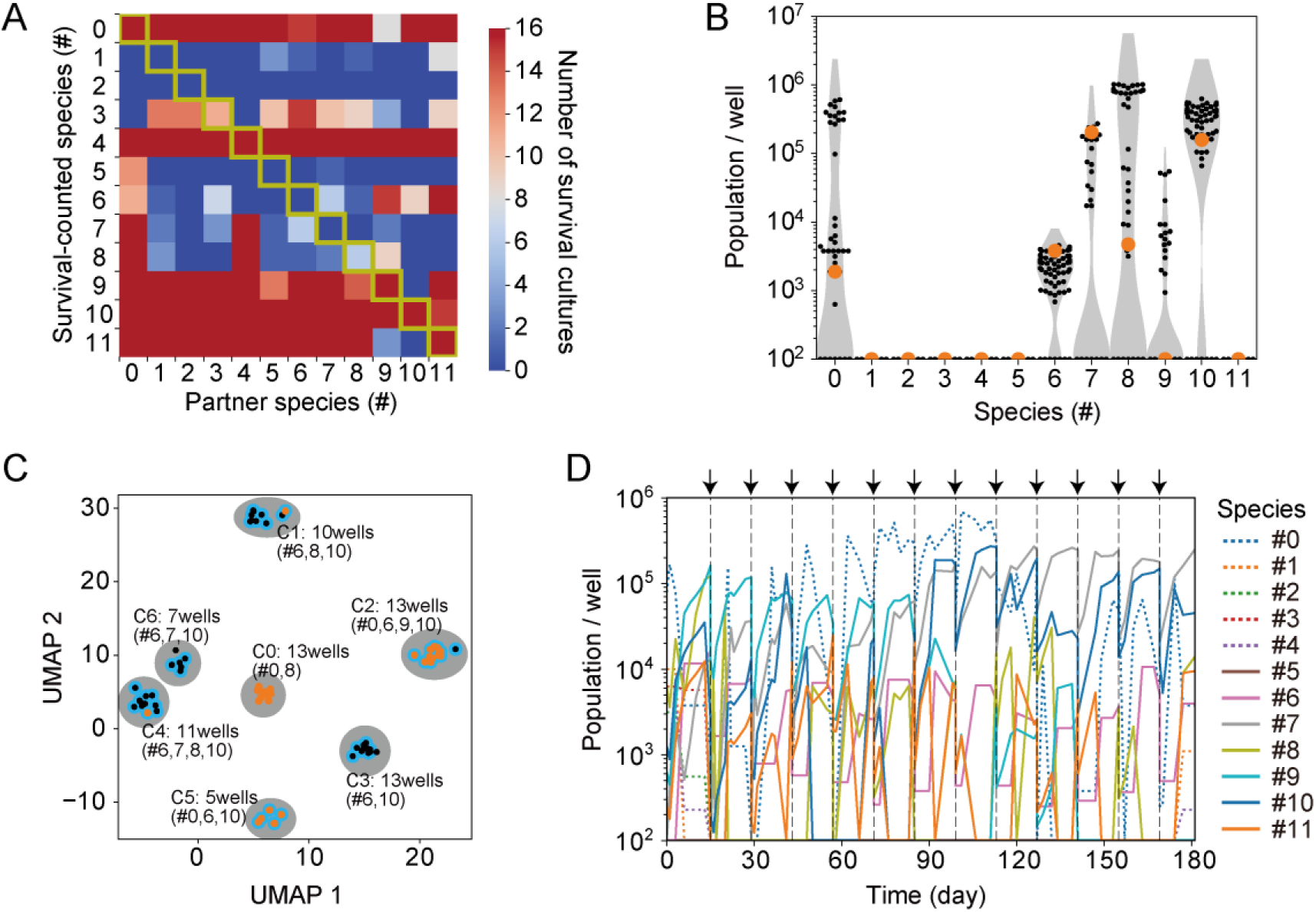
Results of semi-open microcosms. (A) Pairwise cultures. For each pair or monoculture, eight replicates with both the high and low initial concentrations were tested (16 cultures in total; the correlation coefficient between the results with the high and low initial concentrations was 0.9; see Figure S6 for details). The color shows the number of cultures in which the species on the vertical axis survived. The horizontal axis shows the partner species of the species at the vertical axis. For instance, a point with a horizontal axis value of 6 and a vertical axis value of 9 represents the results of co-culture between species #6 and #9, indicating the number of co-cultures with survival of species #9. On the diagonal, a point with a horizontal axis value of 9 and a vertical axis value of 6 corresponds to the same co-culture experiment, but it illustrates the survival of species #6. Pairs of the same species indicate monocultures, depicted as yellow-edged squares. (B) Population of each species in eleven-species microcosms with low initial concentration after six months. Black dots show the data for each of 72 wells, and gray background shows their violin plot. Orange plots show the results of an example well shown in (D). These results after six months were first quantified using machine learning. Subsequently, visual verification was conducted to determine only the presence or absence of each species to avoid the effect of misclassifications. Species that were not observed were recorded as zero. Additionally, in the samples from these six-month cultures, all instances that machine learning identified as species #11 were found to be misclassifications of species #10; thus, the counts for #11 were reassigned to #10. (C) UMAP analysis of the results shown in (B). Each dot shows the result of each well. Orange and black dots show the wells in which *E. coli* was detected and not detected, respectively. Blue-edged dots show the wells in which *T. thermophila* was detected. Gray background circles show the clusters by the k-means method. The results of the wells with the same cluster were similar, and typical survival species are indicated as text in the plot, although some wells have one more or one less species. (D) Example population dynamics. The results of the well depicted as orange circles in (B) are shown. Gray arrows depicted at the top of the graph show the time point of subculturing with 1/10 dilution, conducted approximately once every 2 weeks. The population of each species was determined by the combination of three noninvasive measurements using the fluorescence spectroscopy, machine learning, and the temporal variation of micrographs (see Text S2 for details). Unlike the results for B and C mentioned above, these results have not been corrected through visual inspection and thus include the impact of misclassifications by the machine learning (typically, existence of #11 at the six months; see text for discussion).

### Semi-open microcosms

In nature, an ecosystem has a certain boundary and there are inflows and outflows of components. To imitate these flows under controlled conditions, microcosms in the plate wells were closed with a heat adsorption seal immediately after inoculation, opened approximately once every two weeks, and transferred to new wells at a dilution of 1/10 with fresh medium. These microcosms were designated as “semi-open microcosms.” Herein, we show results related to the search for conditions wherein at least a single species from each of the three functional groups (producers, consumers, and decomposers) and one redundant species survive (designated as “target diversity” hereafter).

In semi-open microcosms, the only species that increase by 10-fold in two weeks can survive against the dilution. Therefore, if an individual is detected after half a year (*i.e.* approximately 10^12^ fold dilution in total), it is likely that the species has survived, rather than the dead body accidentally being left through the dilution. We set the initial concentration of each species to be 1/200 (blue triangles in Fig 2C) or 1/2000 (orange triangles) of the saturating concentration in monocultures (black triangles), and hereafter, we will refer to these initial concentrations as “high initial concentration” and “low initial concentration,” respectively.

We first tested cases wherein two species were inoculated into each well to investigate two-species interactions. Monocultures (single species) under the same experimental conditions were also tested for comparison. Figure 3A shows a summary of the survival results for each species after six months (in this case, the two types of initial concentrations showed similar results; Figure S6). Positive and negative interspecific interactions were observed. For example, neither of the two consumers (#5 and #6) survived on their own, but co-culturing with *E. coli* made them viable (predator-prey relationship). A green alga *R. subcapitata* (#9) survived alone, but did not survive when co-cultured with *C. vulgaris* (#10; competition). In the pair of a consumer *T. thermophila* (#6) and a cyanobacterium *A. circularis* (#7), both species benefitted from each other. Note that since the culture medium contains nutrients, even decomposers can grow on their own. Conversely, because some nutrients are often toxic to cyanobacteria (Hosoda et al., 2014), co-culture with decomposers can enhance their survival.

When the 12 species were mixed, both the consumers (species #5 or 6) were absent after six months in all tested 128 wells (64 replicated wells for high and low initial concentrations, respectively). Instead, only *C. parapsilosis* (#4) survived among the eukaryotic nonproducers. Given that a consumer species survived under conditions wherein *C. parapsilosis* was excluded (as described below), this species might have had a competitive relationship with the consumers, possibly due to direct effects or indirect interactions with other species.

We then tested microcosms with 11 species, excluding *C. parapsilosis*, to identify conditions under which at least one consumer species would survive. When the 11 species were mixed at high initial concentration, a consumer *T. thermophila* (#6) survived after 6 months in 62 of 64 tested wells. In 4 of the 62 wells, *E. coli* (#0), *T. thermophila* (#6), *R. subcapitata* (#9), and *C. vulgaris* (#10) survived, which satisfies the target diversity. However, no prokaryotic producers survived among them.

When the 11 species were mixed at low initial concentration, 15 of the 72 tested wells maintained the target diversity. Figure 3B shows the population of each species after six months. Although six species remained, we did not observe any wells in which all six species survived. In other words, wells that were aliquoted from the same test tube, implying nearly identical initial conditions, showed different surviving species. This probabilistic nature of outcomes was found due to the capabilities of the high-throughput experiments.

Figure 3C summarizes the results of 72 wells after six months using UMAP (Uniform Manifold Approximation and Projection) analysis (McInnes et al., 2018). One point indicates the state of one well after six months. The gray circles represent the seven clusters collected by k-means clustering. First, *C. vulgaris* was alive in all wells except for cluster C0. In addition, *T. thermophila* survived in all but two of these wells (blue edges). Of these, clusters C1, 2, 4, 5, and 6 (*i.e.*, except C0 and C3) had multiple surviving producers. Furthermore, among these, the orange dots showed the target diversity because *E. coli* was present (16 wells). Note that *E. coli* was not detected by the plate reader and microscope in many wells where *T. thermophila* survived; however, we observed the growth of *E. coli* in all wells when seven of these wells were transferred to LB medium, indicating that *E. coli* survived below the detection limit.

Figure 3D shows the population dynamics of one example of the 16 wells with the target diversity (also shown in Figure 3B; orange circle). We confirmed the coexistence of *E. coli* (#0), *T. thermophila* (#6), *A. circularis* (#7), *Synechocystis sp.* PCC 6803 (*S*. 6803; #8), and *C. vulgaris* (#10) in this well, using both machine learning and the eyes. This well contained both prokaryotic and eukaryotic species for both consumers and producers.

The results of the 11-species mixture also revealed interesting phenomena that could not be accounted for by any single two-species interaction. Specifically, cluster C0 contained only *E. coli* (#0) and *S*. 6803 (#8). Although *C. vulgaris* (#10) was able to survive when co-cultured with all other species in the pairwise experiments (Figure 3A), it was absent from this cluster. This observation might be due to the absence of a predator, *T. thermophila* (#6), leading to a high population of its prey, *E. coli*. This elevated *E. coli* population could have provided a competitive advantage to its mutual partner, *S*. 6803, resulting in the exclusion of *C. vulgaris*. In addition, *C. vulgaris* did not coexist with any other producers in the pairwise experiments, but coexisted with the other three producers (#7-9) in the 11-species experiments. These results suggest that other species (*i.e.*, diversity) provided a new competitive niche. As mentioned above, a probabilistic nature and interesting interspecific interactions were observed, and a quantitative analysis of these phenomena is expected in the future.

### Closed microcosms

Closed systems are important for understanding the earth and terraforming, and for clarity as a physical system (Armentrout et al., 2000; Hekstra and Leibler, 2012; Nakajima, 2021; Taub, 1974; Taub and McLaskey, 2014). In closed systems, the wells were sealed at all times using a heat absorption seal. Unlike semi-open systems, proliferation is not guaranteed, even if an individual organism exists; therefore, it is difficult to determine whether a species is alive or dead. It was impossible to measure the population using microscopy due to the high concentration under these conditions (Figure S4A) because organisms were not eliminated regardless of whether they were alive or dead. Thus, fluorescence spectroscopy was used to quantify the population. Despite these problems, we found experimental conditions that maintained the target diversity for at least 6 months.

When 12 species were mixed at low initial concentration, the consumers (species #5 or 6) were absent at 6 months in all 32 tested wells, and only cyanobacterial fluorescence was detected (Figure 4A). Similar results were obtained when the 11 species (except *C. parapsilosis*) were mixed at low initial concentrations.

**Figure 4.**
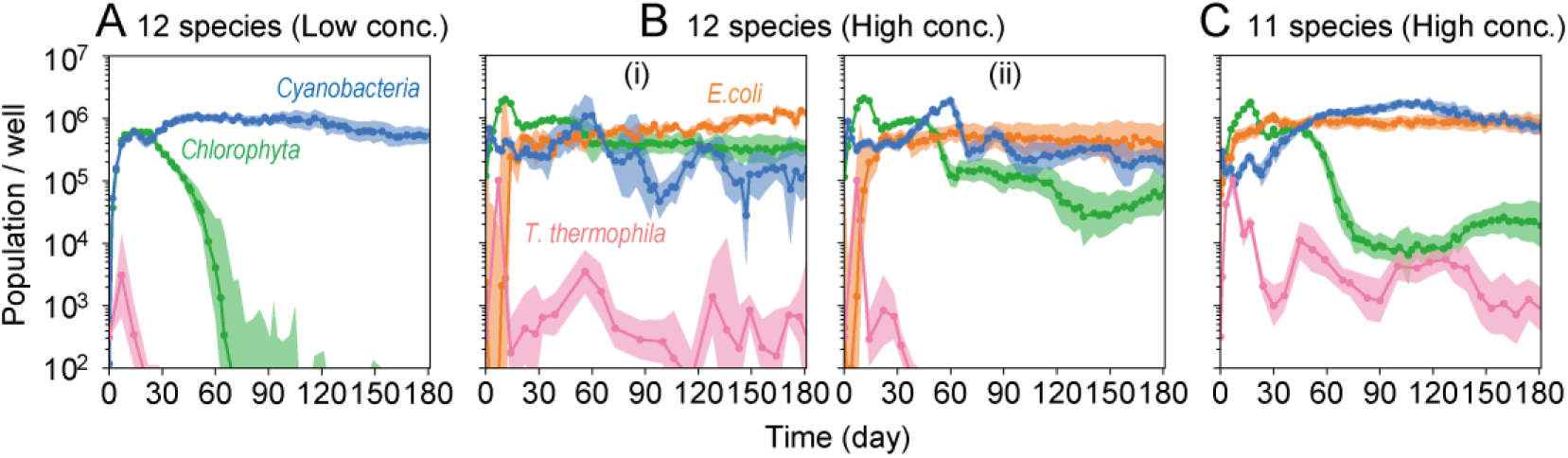
Population dynamics of closed microcosms. The populations of *E. coli*, cyanobacteria, and green algae were quantified by fluorescence spectroscopy. The *T. thermophila* population was quantified from the temporal variation of micrographs. The dots with lines indicate the geometric mean values, and the shaded regions include one geometric standard deviation above and below the geometric mean (population data below 1 was set to 1 for log calculation). (A) 12 species microcosms with low initial concentration (32 wells). (B) 12 species microcosms with high initial concentration. The results are shown separately for the wells in which (i) *T. thermophila* survived for six months (3 wells) and (ii) did not (29 wells). (C) 11 species microcosms with high initial concentration (32 wells).

When the 12 species were mixed at high initial concentration, a consumer (*T. thermophila*) was detected in three of the 32 tested wells. The fluorescence of *E. coli*, cyanobacteria, and green algae was also detected in the three wells (Figure 4B-(i)). Thus, these wells satisfied the target diversity requirements and contained at least one bacterial decomposer (*E. coli*), one eukaryotic consumer (*T. thermophila*), one bacterial producer (cyanobacteria), and one eukaryotic producer (green algae). Using microscopy, immediately after shaking by inversion, we observed the existence of *C. parapsilosis*, *S. 6803*, *R. subcapitata*, *C. vulgaris*, and *C. reinhardtii*, in addition to *E. coli* and *T. thermophila* (thus, at least seven species) in all three wells (Figure S4B), although we could not determine whether they were alive (proliferative). As a more stable condition, in the case of the 11 species microcosms (other than *C. parapsilosis*), at least *E. coli*, *T. thermophila*, one cyanobacterium, and one green alga were maintained with population changes in all 64 wells tested (Figure 4C), thus satisfying the target diversity.

Population dynamics also include the following interesting phenomena. For example, although *T. thermophila* is a predator of *E. coli*, the prey *E. coli* population continued to increase and decrease after 60 d in the presence (Figure 4B-(i)) and absence (Figure 4B-(ii)) of the predator, respectively, in the 12-species microcosm (both significant slopes were detected by the regression *t*-test, α=0.05). No significant increase or decrease in *E. coli* after 60 d was observed in eleven-species microcosm (Figure 4C). In addition, the concentration of *E. coli* after 6 months was significantly lower only in the absence of *T. thermophila* (Figure 4B-(ii)) than in the 11 (Figure 4C) and 12 (Figure 4B-(i)) species microcosms in the presence of *T. thermophila* (Tukey’s HSD, α=0.05). The fact that the prey population was larger in the presence of the predator suggests that there was a higher-order interaction.

In addition, stepwise transitions were consistently observed (indicated by the green lines in Figures 4B and 4C). These transitions included an initial rise during the first 10 days, a slight decline, stabilization between days 20 to 40, and a subsequent decrease. These observations highlight the intriguing dynamics inherent to this ecosystem. These constrained transitions align with the concept of homeorhesis in ecological succession (Chuang et al., 2019; Odum and Barrett, 2005). Future studies will focus on the quantitative analysis of these interesting observations.

## Conclusions

In conclusion, we developed a model ecosystem with 12 diverse axenically culturable and cryopreservable microbial species. We propose a method using machine learning for the noninvasive quantification of these species, which allows high-throughput experiments. We found experimental conditions that allowed the maintenance of at least one species from each of the three functional groups (producers, consumers, and decomposers) and one redundant species for six months both in semi-open and closed systems. These conditions serve as scaffolds for future studies. Although not all 12 microbial species coexisted in this study, future endeavors to explore varying experimental conditions or the addition of species, aiming for greater coexistence, would further our understanding of ecosystem dynamics and mechanisms of coexistence. We also found interesting phenomena, such as probabilistic consequences, high-order interactions, and stepwise transitions, which are expected to be analyzed in detail in the future. Our model ecosystem is suitable for experimentally investigating evolution, such as massively conducted studies at the individual level (Blount et al., 2018), and for understanding species interactions, which require high-throughput experimental systems for a vast number of species combinations. Therefore, our model ecosystem will accelerate experimental approaches to study evolutionary and ecological dynamics.

## Materials and Methods

### Strains

The *E. coli*strain DH1ΔilvE::(dsred.T3-cat) was constructed previously (Hosoda et al., 2011). *S. epidermidis* strain JCM2414 and *C. parapsilosis* strain JCM1612 were provided by the Japan Collection of Microorganisms (JCM), RIKEN BRC, Japan. *S. pombe* strain L972 was a kind gift from Dr. Kojiro Ishii of the Kochi University of Technology. *S. cerevisiae* strain BY25611 (BY4741ho::Kan-TEFIIpr-GFP-ADH1ter) (Breslow et al., 2008) was provided by the National Bio-Resource Project (NBRP), Japan. *D. discoideum* strain AX2 (DBS0235521) was provided by the Dicty Stock Center, Northwestern University, USA. *T. thermophila* was a conjugation progeny of strains CU427 and CU428 (to observe evolutionary changes in future studies), provided by the Tetrahymena Stock Center, Cornell University. *A. circularis* strain NIES-21, *R. subcapitata* strain NIES-35, *C. vulgaris* strain NIES-227, and *C. reinhardtii* strain NIES-2236 were obtained from the NIES collection, Japan. *S*. 6803 (ATCC27184) was obtained from the American Type Culture Collection (ATCC, USA).

### Monocultures

*E. coli* was cultured in LB medium (Bertani, 1951) at 23°C. S. epidermidis, S. pombe, S. cerevisiae, and C. parapsilosis were cultured in YPD medium (Sherman, 2002) at 23°C. D. discoideum was cultured in HL5 medium (Cocucci and Sussman, 1970) at 22°C. *T. thermophila* was cultured in modified Neff’s medium (Cassidy-Hanley, 2012) at 23°C. A. circularis was cultured in BG11 medium (Allen, 1968) at 20°C with irradiation by a white LED panel (TH-224X170SW; CCS, Japan) at an intensity of 8 μmol·m^−2^·s^−2^ for 12 h intervals. S. 6803, R. subcapitata, C. vulgaris, and C. reinhardtii were cultured in BG11 medium at 23°C with irradiation at an intensity of 50 μmol·m^−2^·s^−2^ for 12 h intervals.

### Microcosms

For microcosm experiments, we prepared 50 µL of culture solution in each well of a 384-well plate (#142761; Nunc, USA) and sealed the plate with a heat adsorption seal (#4ti-05481; 4titude, UK) (the input concentration of microorganisms is described at each part of the results). As the plate and seal were made of a polymer, they were not completely sealed. The edge and four corner wells were dried for approximately 40 and 20 months (approximately 1μL/month per side). We did not measure the concentrations of gas and dissolved gas in the wells; however, carbon dioxide was suggested to be deficient because the saturation concentration of phytoplankton in the sealed well was lower than that in the unsealed monocultures (approximately two orders; Figure 2C).

We used liquid medium BG11HLB, which contains half the concentration of BG-11 medium for cyanobacteria (Allen, 1968) and 1/100 the concentration of LB medium for bacteria (Bertani, 1951). We added the microorganisms to the medium in each well of a 384-well plate and placed the plate on a white LED panel (TH-224X170SW; CCS, Japan) in an incubator at 23°C with irradiation at an intensity of 50 μmol·m^−2^·s^−2^ for 12 h intervals. The plate was shaken by inversion and spun down twice a week; otherwise, the ceiling water continued to increase. Note that the generation of ceiling water was prevented when light was illuminated from the top; however, the results of multiple species coexistence shown in this study were not obtained using the same black 384-well plate.

### Measurements

A fluorescence plate reader (Varioskan Flash; Thermo, USA) was used for fluorescence spectroscopy to quantify the concentrations of cyanobacteria, green algae, and *E. coli* (red fluorescent protein-labeled) from the corresponding fluorescence (Table 1). Before measurement, the plate was shaken by inversion and then spun down. Calibration curves for each species and the separation between the cyanobacteria and green algae are shown in Figure S7.

For microscopy, the plate was left for 3 h after shaking and then spun down for fluorescence spectroscopy before observation. We scanned each well using an inverted microscope (Nikon Ti-E with the Perfect Focus System and High Content Analysis; Nikon, Japan), and obtained bright-field images (two images with a time difference of 12 s), and fluorescent-field micrographs (three images with filter sets of Semrock FITC-3540C, Semrock TRITC-B, and Chroma 49006-ET-Cy5) of the center of each well (one position per well) from the bottom. 60X objective lens (CFI S Plan Fluor ELWD ADL 60XC, Nikon, Japan) or 4X objective lens (CFI Plan Apochromat Lambda D 4X, Nikon, Japan), and a digital CMOS camera (Neo sCMOS, Andor, UK; 2048 × 2048 pixels; 0.1 µm/pixel) was used for capturing images. We did not apply phase-contrast observation, but bright-field observation for transmitted light microscopy. The wells of the 384-well plate were too narrow to be adopted for phase-contrast observation, and even small particles such as bacteria could be seen with bright-field observation because the light reflected off the wall of the narrow wells.

For quantification of the concentration of *T. thermophila* (a consumer), the temporal variation of images obtained by low-magnification micrographs using the 4X objective lens, whose viewfield (one side is approximately 3.3 mm with a resolution of 1.6 µm/pixel) captured the entire well, because there was a correlation between *T. thermophila* concentration and the temporal variation since *T. thermophila* is large (major axis approximately 50 µm) and swims. Specifically, we utilized the two bright-field images mentioned above, taken with a 12-second interval. For each image, the luminance values were linearly converted such that the quantiles of 0.2 and 0.8 were rescaled to the values of 0 and 1, respectively. The images were subsequently resized to 512 × 512 pixels. The absolute differences between each pixel in the two images were then computed. We counted the number of pixels for which the absolute difference exceeded a value of 0.4. The calibration curve for this value is shown in Figure S8.

### Machine learning methods

To build a model for detecting individual microorganisms in microcosm images obtained using the 60X objective lens, a training dataset for supervised learning was annotated using a combination of classical image processing methods. Wells with only a single species were prepared and microbial detection was performed using combinations of preprocessing, filters, and segmentation algorithms. Owing to the distinct shapes (rod, oval, and filamentous) and properties (signals in bright-field or fluorescence) of each species, we tailored our methods to extract regions representing individual cells or multi-cell clusters from microscopic images based on the specific species type. Although not all individuals were detected, thousands of bounding boxes (locations of individual microorganisms) were obtained. The input channels and combinations of algorithms for extracting the regions are listed in Table S1, and examples of the segmentation results are shown in Figure S1.

For the machine learning model, we used the publicly available object detection network framework YOLOv3 (Redmon and Farhadi, 2018) and performed retraining using the dataset to detect and classify the microorganisms. The code used for YOLOv3 was cloned from the keras-yolo3 (qqwweee, 2023). The network weights were downloaded from the YOLO website (Redmon) and converted into a Keras model. Bright-field and three fluorescent-field images of each well were incorporated into a single RGB image (bright-field into all RGB channels and Semrock TRITC-B, Chroma 49006-ET-Cy5, and Semrock FITC-3540C into R, G, and B, respectively) at an intensity ratio of 7:3 (bright-field: fluorescent-field). The Images were cropped to 512 × 512 pixels from the original 2048 × 2048 pixels and the information of the bounding boxes was set appropriately. We performed retraining for 100 epochs using Adam optimizer and standard data augmentation (batch size = 32; learning rate = 10^−3^ for the first 50 epochs, then 10^−4^). For each species, an average of 7.5 microscope images (SD = 1.9) was used as training data, with an average individual count of 685.6 (SD = 503.7). This underscores the efficacy of transfer learning, which can yield a good performance even with a limited number of training images. Moreover, in a more general case, when adding a new class (in this context, a new species), even simple deterministic operations conducted in a non-parametric manner are known to achieve accuracy levels of approximately 80% for existing classes using only a single training instance (Hosoda et al., 2022).

We detected individual microorganisms in images of microcosms or standard monoculture samples and used the detected data with YOLO confidence scores (Redmon et al., 2016) of 0.5 or higher. This machine learning model recognizes species based on the shape and fluorescence of each individual microorganism and does not determine whether the individual is alive or dead.

### Population quantification

We measured the microcosms using the fluorescence plate reader approximately once every 3 or 4 days, and before and after subculturing on the same day. In addition, micrographs were obtained approximately once every one or two weeks. Micrographs were required to obtain the population of *T. thermophila* from the temporal variation in the image (see above) and to obtain the populations of other species using the machine learning model. We combined the data from the plate reader and the micrographs to obtain the population dynamics at the time resolution of the plate reader as follows.

We reconstructed the microscopy data by applying the most recent microscopy data each time a measurement under the microscope was missing (summarized in Figure S9A). The value after subculturing was set to 1/10 of the value before subculturing, according to the experimental procedure. These reconstructed values were directly used as the population for non-fluorescent species (#1–6; Figure S9B; including *S. cerevisiae*, which was GFP-labeled, but not detected by the plate reader because the fluorescence was too weak). The population of *E. coli* can be measured directly using either the plate reader or the machine learning model. The measurement using the machine learning model has greater sensitivity, but is not applicable to dense microcosms (giving a value lower than the truth). Therefore, we applied a larger value of the population determined by the plate reader or the machine learning model to use the value measured by machine learning for the part that the plate reader could not detect. For cyanobacteria and green algae, the total population was measured using the plate reader, and the species composition among each of them was determined by the population ratio using the machine learning model.

## Acknowledgments

We would like to thank Dr. Chikara Furusawa, Kojiro Ishii, and Toshiyuki Nakajima for their valuable advice, and Dr. Tatsuya Nakamura and Mxes. Yoshinori Hiratani, Miwa Kobayashi, Saki Shigeyama, Miki Shimomoto, and Joshua C. Triyonoputro for their technical assistance. We thank Editage for editing this manuscript for English language. We are also grateful to Dr. Isamu Taguchi and the other 108 supporters for the financial support provided through Academist crowdfunding #150. This research was supported in part by JSPS KAKENHI Grant Numbers JP18H04821, JP20H05533, JP20H04868, and JP20K06825, and the “Program for Leading Graduate Schools” of the ministry of education, culture, sports, science and technology (MEXT) Japan. The authors declare no conflict of interest.

## Supplementary material

**Table S1.** Segmentation methods for each species.

|  | Species | Method |
| --- | --- | --- |
| #0 | *Escherichia coli* | Edge detection (DoG filter), thresholding (Otsu), morphology (dilation, fill holes) |
| #1 | *Staphylococcus epidermidis* | Edge detection (DoG filter), thresholding (Otsu), morphology (dilation, fill holes) |
| #2 | *Schizosaccharomyces pombe* | Edge detection (DoG filter), thresholding (Otsu), morphology (dilation, fill holes) |
| #3 | *Saccharomyces cerevisiae* | Circle detection (Canny filter, Hough gradient method), watershed segmentation |
| #4 | *Candida parapsilosis* | Circle detection (Canny filter, Hough gradient method), watershed segmentation |
| #5 | *Dictyostelium discoideum* | Edge detection (DoG filter), thresholding (Otsu), morphology (dilation, fill holes) |
| #6 | *Tetrahymena thermophila* | Thresholding (local thresholding), morphology (opening, closing, fill holes), watershed segmentation |
| #7 | *Anabaenopsis circularis* | Frangi filter, thresholding (Otsu), morphology (closing) |
| #8 | *Synechocystis* sp. PCC 6803 | Circle detection (Canny filter, Hough gradient method), watershed segmentation |
| #9 | *Raphidocelis subcapitata* | Thresholding (Otsu), morphology (opening, closing) |
| #10 | *Chlorella vulgaris* | Circle detection (Canny filter, Hough gradient method), watershed segmentation |
| #11 | *Chlamydomonas reinhardtii* | Circle detection (Canny filter, Hough gradient method), watershed segmentation |

**Figure S1.**
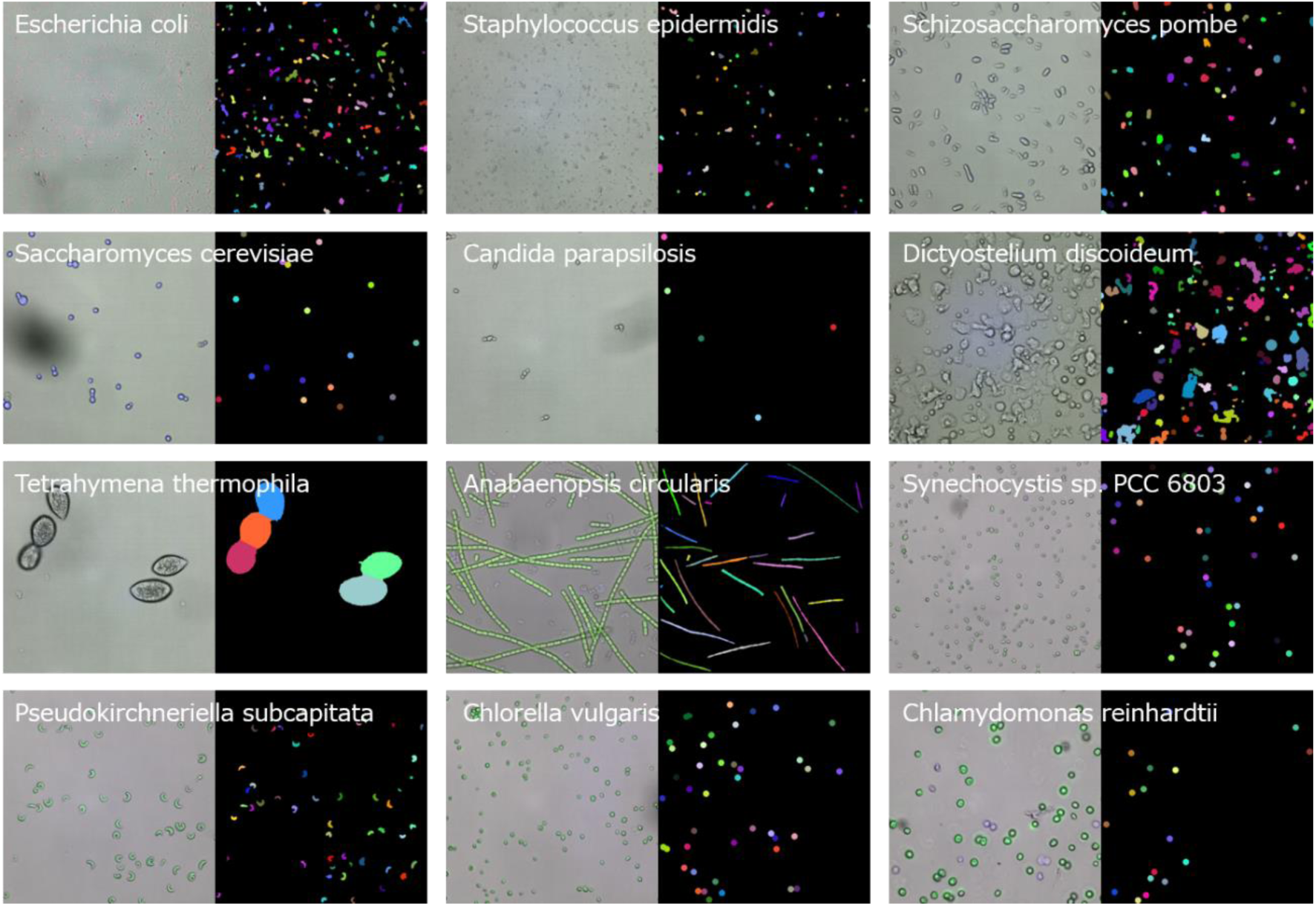
Segmentation results by classical methods. For each species, the left side is a representative micrograph of bright-field and fluorescent-field superimposed (one side is approximately 222 μm; see Text S1 for RGB colors). The right side shows segmentation into each individual. The mask of each individual is represented by the same color.

**Figure S2.**
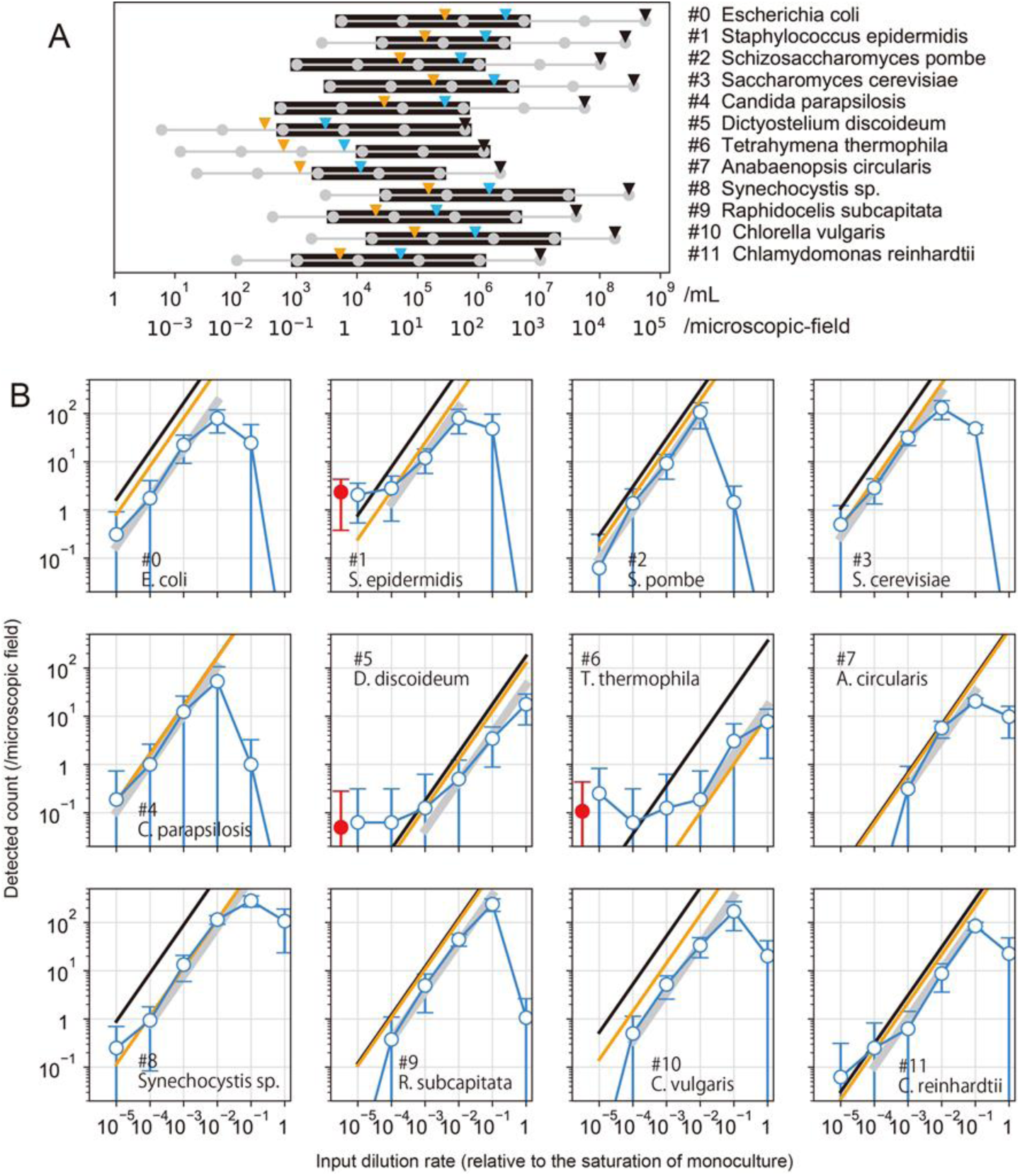
Standard curves of measurements using the machine learning model for each species. The horizontal axes show the dilution rate from the saturated monocultures. Blue circles with lines show the mean detected count in each micrograph of 16 replicated wells from 4 independent plates. Red circles show the mean detected count of 384 blank wells (only medium). The error bars show the standard deviation. We also counted the number of individuals in each micrograph with eyes at the appropriate dilution rate for counting (10–100 individuals in each micrograph). The orange lines show the values counted with the eyes, extended to all dilution rates. The black lines show the expected values calculated from the input concentration if all input individuals are assumed to sink to the bottom of the well and are evenly distributed on the bottom. Thus, the difference between the orange and black lines shows the fraction of individuals that sunk to the bottom and entered the microscopic view, which is independent of the detection using the machine learning. We determined the concentration range where the counted amount correlates with the input amount as given below. For each species, we fitted the blue circle data with a straight line with a slope of 1 in a log-log plot. We determined the range in which the residuals of all the points were smaller than half order. Gray bold lines show the fitted line with the determined range.

**Figure S3.**
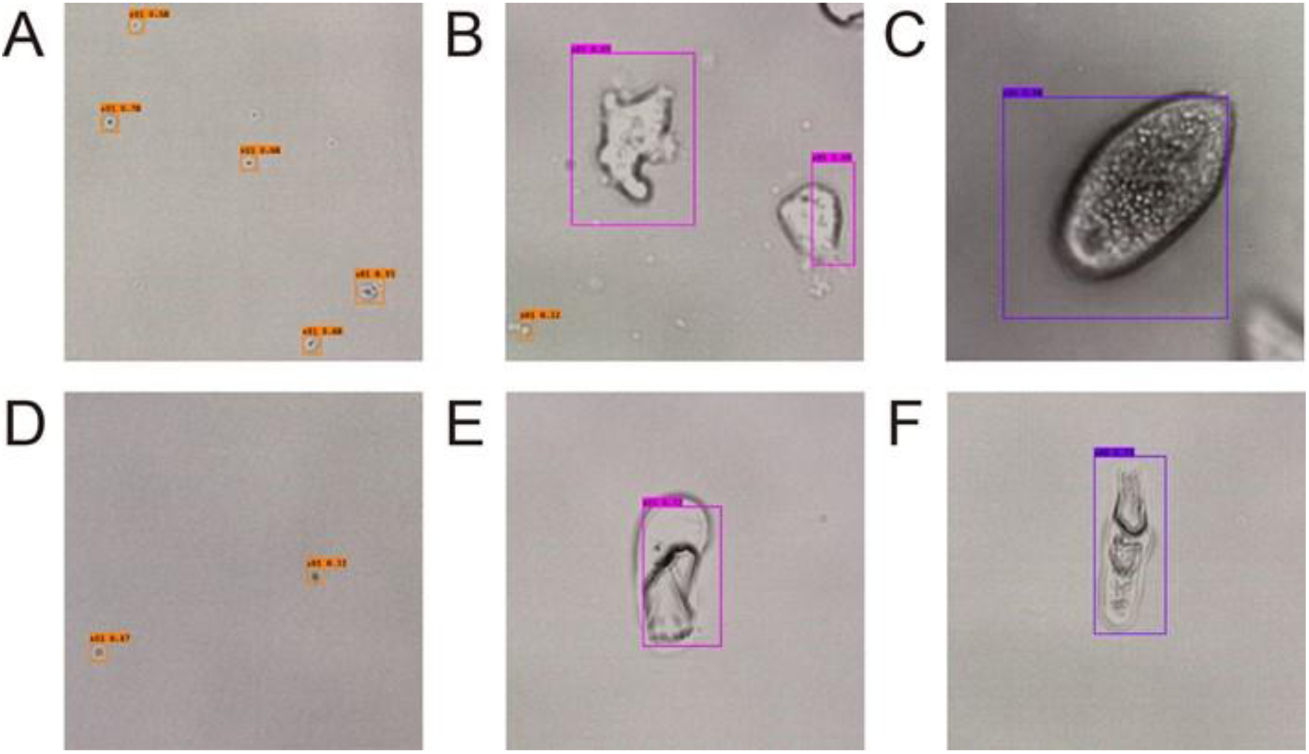
Images of objects detected in blank samples. Our machine learning model detected some objects as species #1, #5, and #6 even in the blank samples, as shown in Figure S2 (red circles). A, B, and C are examples of species #1, #5, and #6, respectively. D, E, and F are examples of objects detected as species #1, #5, and #6, respectively, in the blank samples. One side is approximately 70 μm in all images. Colored orange, magenta, and purple boxes indicate the detection of #1, #5, and #6, respectively. The objects detected in the blank were not due to the medium (for example, scratches on the bottom surface) because they were observed even in wells that contained only ultrapure water. In B, we can also observe the false detection of tiny particles from protozoan #5 as cocci #1.

**Figure S4.**
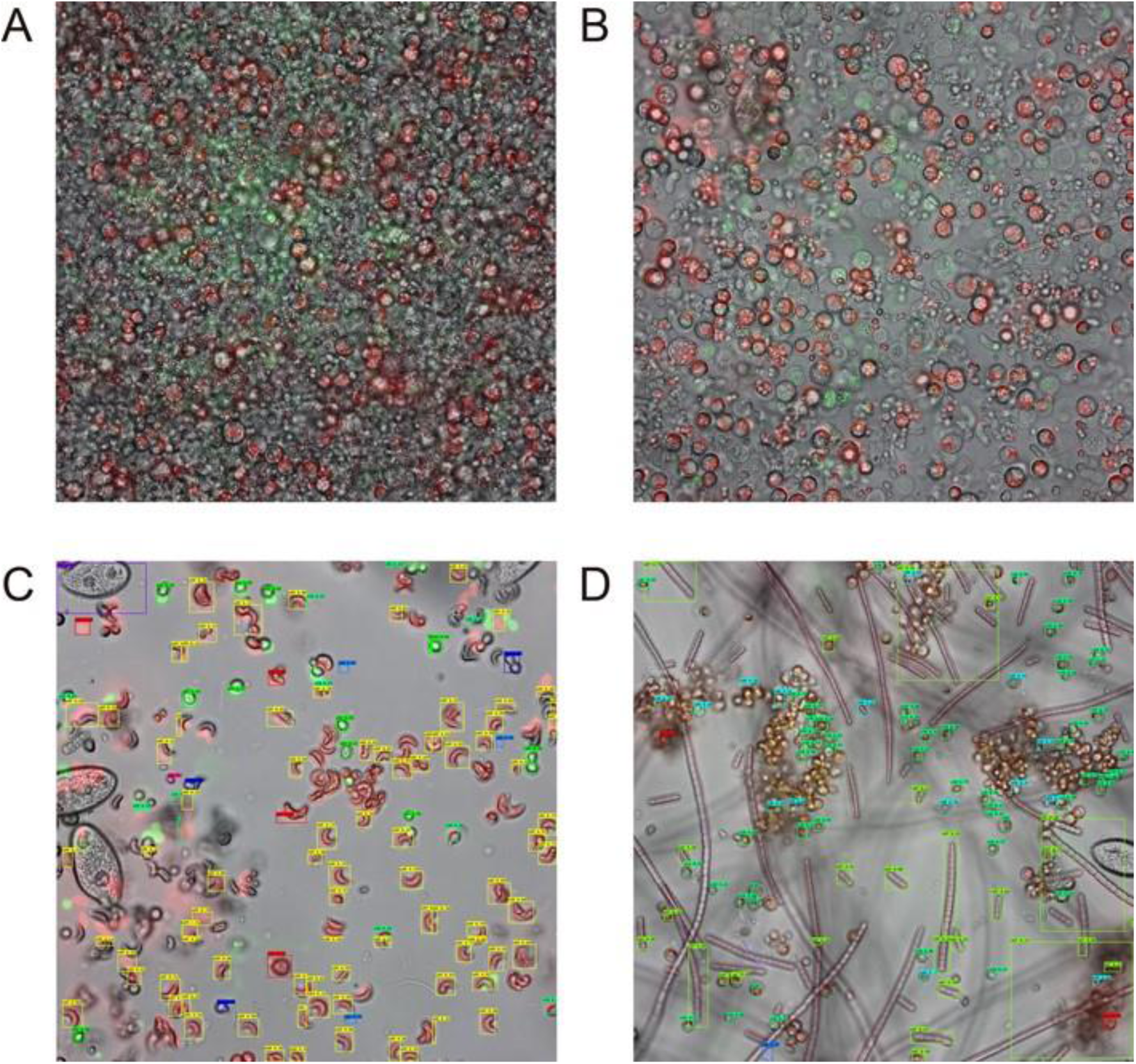
Examples of micrographs of microcosms (one side is approximately 222 μm; see Text S1 for RGB colors). (A) A micrograph of the well of closed-system microcosm shown in Figure 4B-(i) at 7 months. (B) A micrograph of the same well immediately (approximately 10 min) after shaking by inversion. (C) A micrograph of the well of semi-open-system microcosm shown in Figure 3D at 1 week. The positional misalignment between fluorescence and bright fields is due to the movement of the ciliate cells during those time lags. Color boxes show the results of the machine learning detection. (D) A micrograph and the detection results of the same well at 6 months. We observed some clusters, but these clusters floated when shaken and we did not observe biofilms that stuck to the bottom.

**Figure S5.**
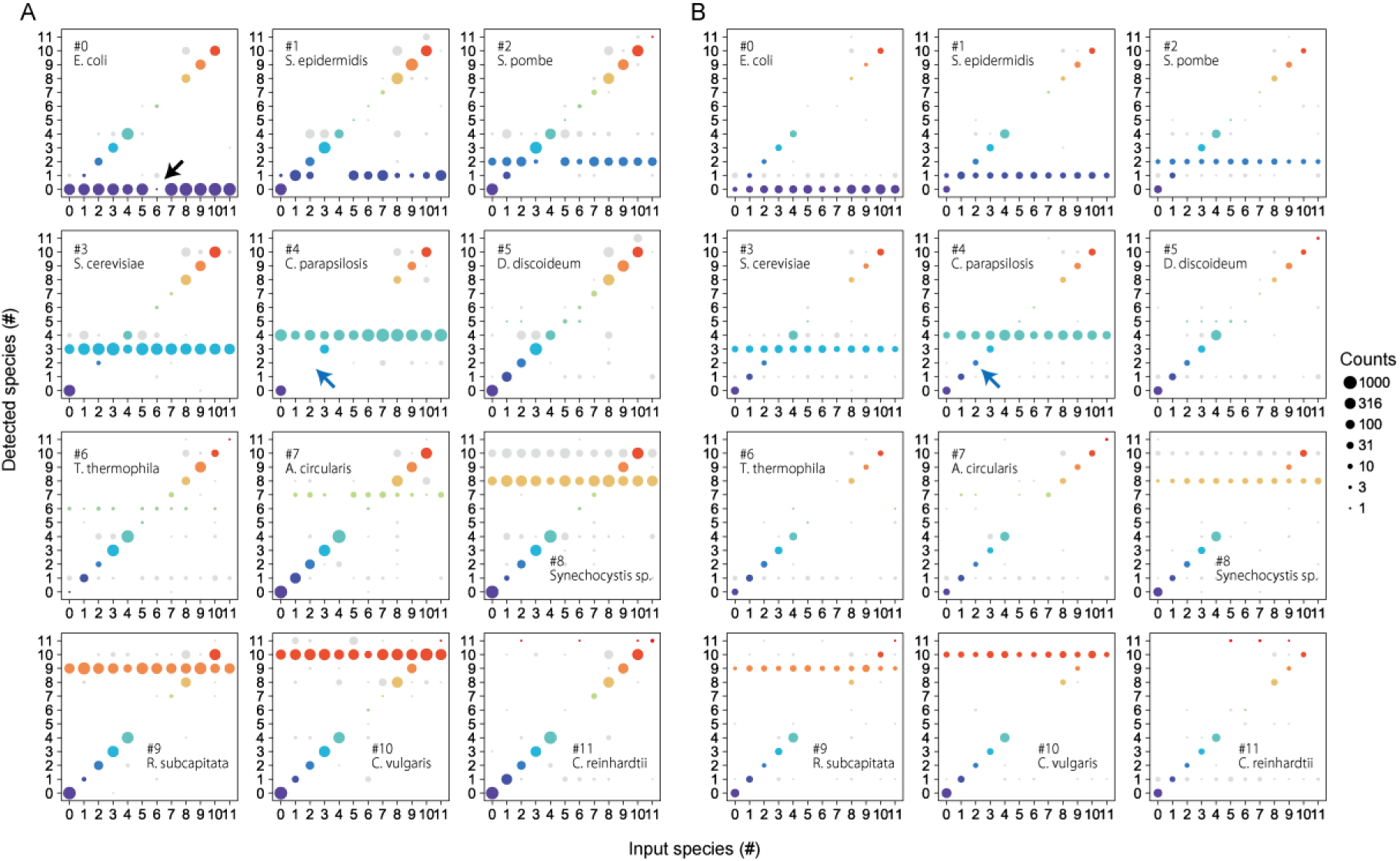
Interference by another species in detection using machine learning. This figure presents the detection results from machine learning when two species are mixed. Panels A and B display outcomes when two species were mixed at 1/200 and 1/2000 of their saturation concentrations, respectively, in monoculture. In each plot, the mixed pair consists of the species denoted by the text within the plot and the species represented on the x-axis. The y-axis indicates the detected species. As an example, examine the plot in the second row and second column of panel A. All experiments in this plot involved species #4, as indicated in the text. When the x-axis was zero, it represented a combination of species #0 and #4. The counts at positions 0 and 4 on the y-axis denote the detection of species #0 and #4, respectively. However, the absence of species #2 in combination with species #4 (highlighted by a blue arrow) serves as an example of detection inhibition due to high concentrations (as observed, for instance, in Figure S4A). While the non-detection of #2 was pronounced, there should be a decrease in the counts of #4 from the expected numbers. Panel B illustrates the results when both concentrations were scaled down to 1/10. Even at this 10-fold dilution, species #2 remained detectable when mixed with species #4 (highlighted by a blue arrow). In principle, machine learning detection operates irrespective of whether the sample is a monoculture or mixed when individual organisms are spatially distinct, even within the same microscopic image, given the model design for the 12 classifications. Nonetheless, the spatial overlap within an image can lead to such interference. Although diminished counts in the majority of plots typically indicate interference effects, some plots show predation effects. For instance, the plot at the top left of panel A, with an x-axis value of 6, denotes a mix of species #0 and #6. Comparison of the counts of species #0 when mixed with other species revealed a significant decrease exclusively when combined with #6, suggesting that species #6 rapidly predated species #0 immediately after mixing.

**Figure S6.**
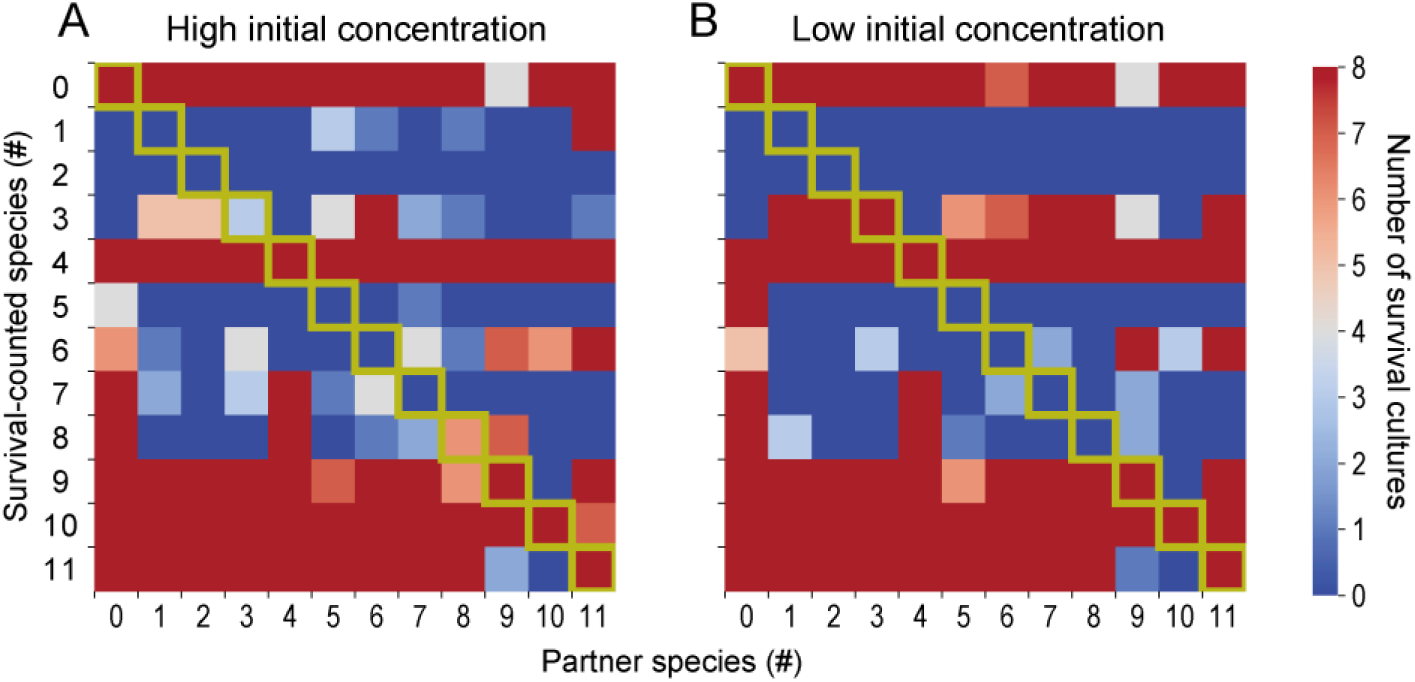
Two-species co-cultures and monocultures in the semi-open microcosms. Results at high initial concentration. (A) and low initial concentration (B) prior to the creation of Figure 3A in the main text. For detailed explanations, please refer to the main body of the paper. Each cultivation result was based on eight replicate experiments. The correlation coefficient between the outcomes of high and low initial concentrations was 0.9.

**Figure S7.**
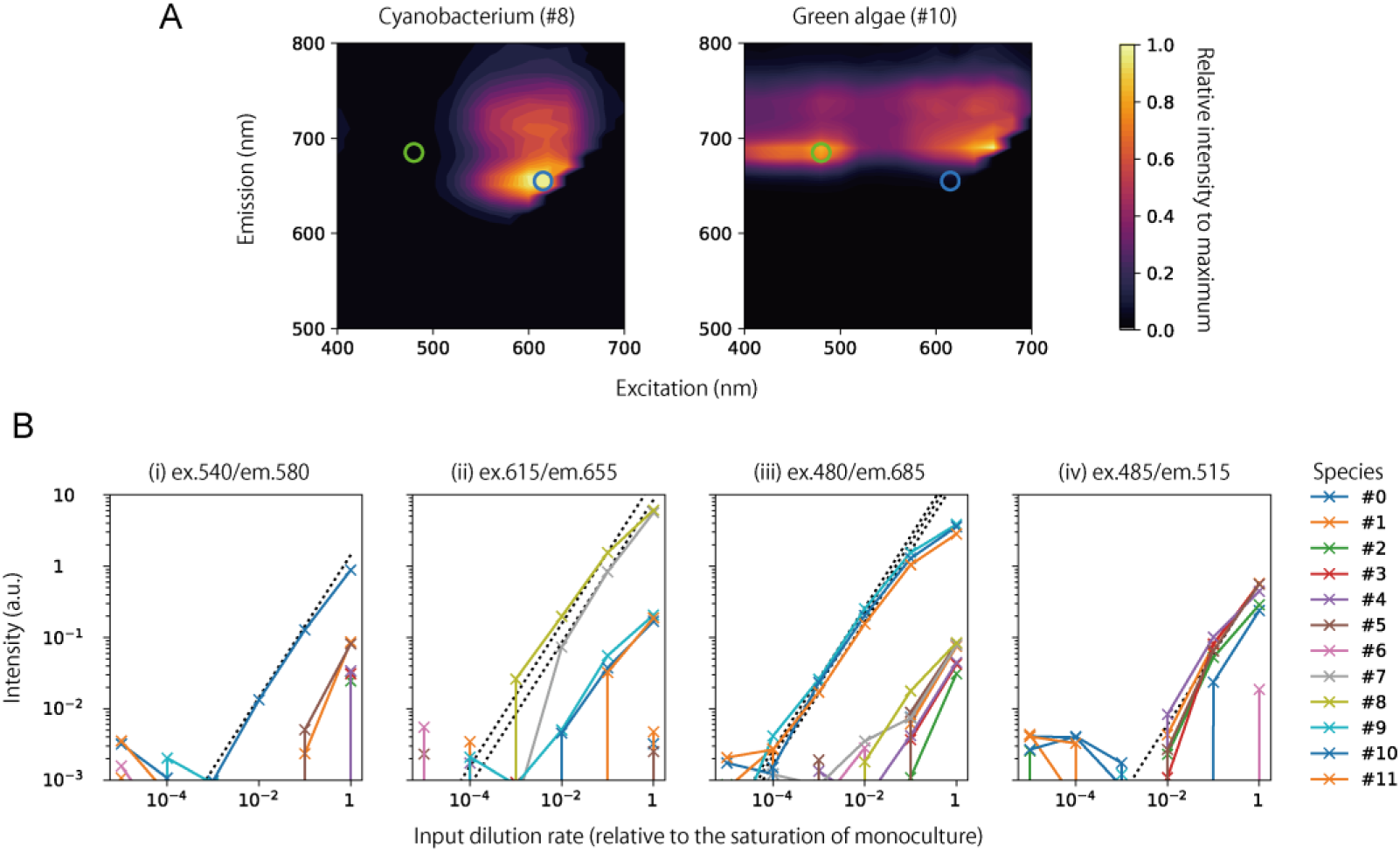
Measurement of each species using a fluorescence plate reader. (A) Distinct fluorescence properties between cyanobacteria and green algae. Species #8 and #10 were used as representatives for cyanobacteria and green algae, respectively. The blue and green circles denote the wavelengths used for quantifying cyanobacteria and green algae in this study. (B) Calibration curves associated with each fluorescence wavelength. All 12 species were individually added at different concentrations into the wells of a 384-well plate. 12 replicate experiments were conducted for each concentration of every species. The figures present the mean value from the 12 replicate experiments, after subtracting the average value from wells containing only the medium, which acted as a background. The black dashed line indicates the calibration curve utilized in this study. (i) The fluorescence wavelength for quantifying RFP-labeled *Escherichia coli* (#0). At the highest concentrations, interferences from other species became apparent. However, such high concentrations never occurred in the microcosm experiments and, thus were considered as negligible. (ii) The fluorescence wavelength for detecting cyanobacteria, corresponds to the blue circles mentioned in (A). Interference from green algae was noted, and it was 7% of the fluorescence intensity of the wavelength for green algae on average. Therefore, the measured intensity was adjusted by subtracting 7% of the fluorescence intensity of the wavelength for green algae in this study. (iii) The fluorescence wavelength for quantifying green algae, aligns with the green circle in A above. Interference from cyanobacteria was observed, which was 1% of the intensity of the wavelength for cyanobacteria on average. Hence, the observed intensity was adjusted by subtracting 1% of the intensity of the wavelength for cyanobacteria. (iv) The fluorescence wavelength for quantifying GFP-labeled *Saccharomyces cerevisiae* (#3). Unfortunately, its fluorescence was weak and hardly distinguishable from the autofluorescence of several other species. Nevertheless, its greenish hue under microscopic observation has facilitated its detection via machine learning. To date, no conducive conditions have been identified where #3 survives in either 11– or 12-species microcosm experiments. In the future, studies should focus on transitioning to a strain with a greater fluorescence intensity and probing for favorable conditions, aiming to foster a more effective experimental system.

**Figure S8.**
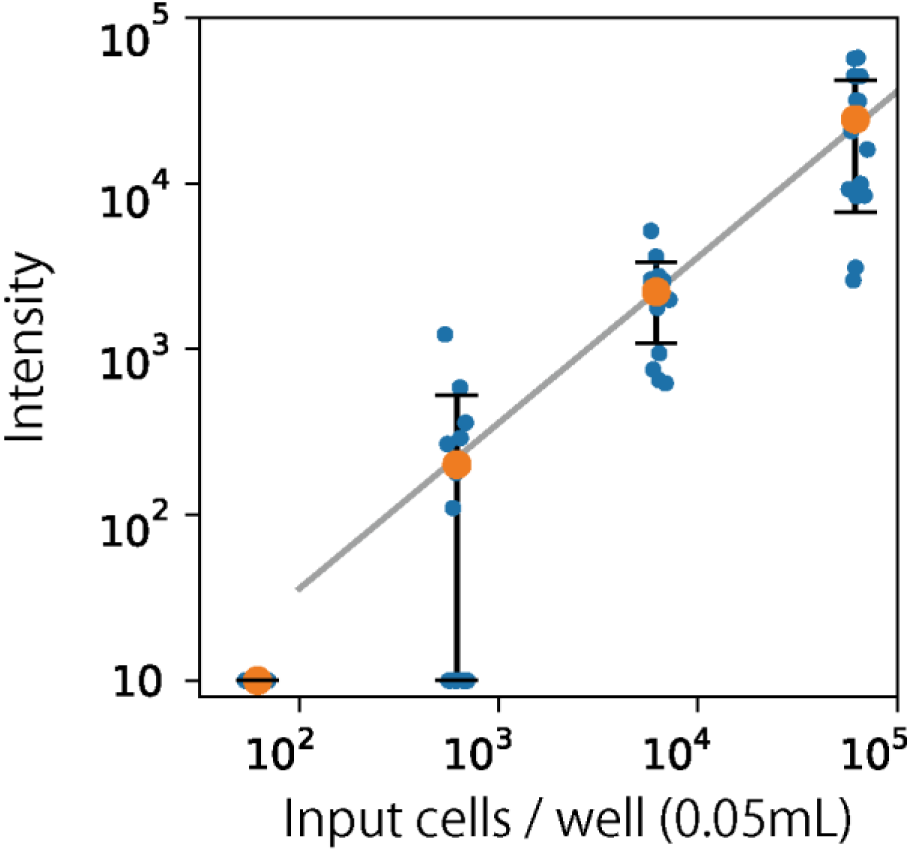
Detection of *Tetrahymena thermophila* based on time-lapse differences in microscopic images. Cells were added to the wells of a 384-well plate at varying concentrations. Sixteen replicate experiments were conducted for each concentration. The blue dots in the figure represent the image difference values for the sixteen replicates, as determined by the method described in the main text. The orange dots depict the average values of the sixteen replicates, with error bars indicating the standard deviation in linear scale. The gray line represents the calibration curve used in this study. It should be noted that *T. thermophila* used in this experiment was taken immediately after monoculture growth. They appeared to be in a nutrient-rich state, showing vigorous movement. In contrast, the microcosm experiments seem to present a more nutrient-deprived environment. Hence, there might be a higher proportion of the cells present on the observable bottom surface in the microcosm experiments. The influence of the biological state on these observations remains an unresolved issue.

**Figure S9.**
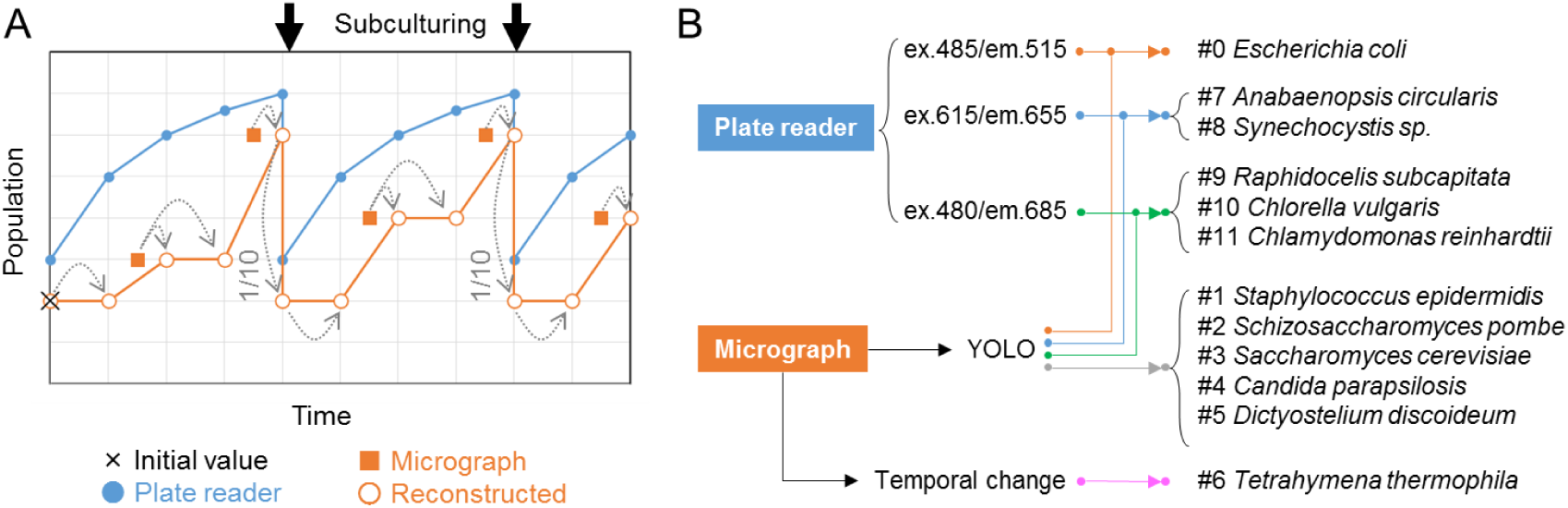
Detailed information about quantifying the population of the 12 species. See main text for explanation.

